# Projecting the end of the Zika virus epidemic in Latin America: a modelling analysis

**DOI:** 10.1101/323915

**Authors:** Kathleen M O’Reilly, Rachel Lowe, W John Edmunds, Philippe Mayaud, Adam Kucharski, Rosalind M Eggo, Sebastian Funk, Deepit Bhatia, Kamran Khan, Moritz U Kraemar, Annelies Wilder-Smith, Laura C Rodrigues, Patricia Brasil, Eduardo Massad, Thomas Jaenisch, Simon Cauchemez, Oliver J Brady, Laith Yakob

## Abstract

**Background** Zika virus (ZIKV) emerged in Latin America & the Caribbean (LAC) region in 2013, and has had serious implications for population health in the region. In 2016, the World Health Organization declared the ZIKV outbreak a Public Health Emergency of International Concern following a cluster of associated neurological disorders and neonatal malformations. In 2017, Zika cases declined, but future incidence in LAC remains uncertain due to gaps in our understanding, considerable variation in surveillance and a lack of a comprehensive collation of data from affected countries.

**Methods** Our analysis combines information on confirmed and suspected Zika cases across LAC countries and a spatio-temporal dynamic transmission model for ZIKV infection to determine key transmission parameters and projected incidence in 91 major cities within 35 countries. Seasonality was determined by spatio-temporal estimates of *Aedes aegypti* vector capacity. We used country and state-level data from 2015 to mid-2017 to infer key model parameters, country-specific disease reporting rates, and the 2018 projected incidence. A 10-fold cross-validation approach was used to validate parameter estimates to out-of-sample epidemic trajectories.

**Results** There was limited transmission in 2015, but in 2016 and 2017 there was sufficient opportunity for wide-spread ZIKV transmission in most cities, resulting in the depletion of susceptible individuals. We predict that the highest number of cases in 2018 within some Brazilian States (Sao Paulo and Rio de Janeiro), Colombia and French Guiana, but the estimated number of cases were no more than a few hundred. Model estimates of the timing of the peak in incidence were correlated (p<0.05) with the reported peak in incidence. The reporting rate varied across countries, with lower reporting rates for those with only confirmed cases compared to those who reported both confirmed and suspected cases.

**Conclusions** The findings suggest that the ZIKV epidemic is by and large over, with incidence projected to be low in most cities in LAC in 2018. Local low levels of transmission are probable but the estimated rate of infection suggests that most cities have a population with high levels of herd immunity.

## Introduction

Starting as early as 2013,^1,2^ the Zika virus (ZIKV) invaded northeast Brazil and began to spread in the Latin America & Caribbean (LAC) region. The subsequent discovery of a cluster of Guillain-Barré syndrome cases and the emergence of severe birth defects led the World Health Organisation (WHO) to declare the outbreak a Public Health Emergency of International Concern in early 2016. The virus has since spread to 49 countries and territories across the Americas where autochthonous transmission has been confirmed.^3^

However, 2017 saw a marked decline in reported Zika cases and its severe disease manifestations.^4^ This decline has been widely attributed to the build-up of immunity against ZIKV in the wider human population,^5^ although it remains unknown how many people have been infected. To date, there has been limited use of population-based surveys to determine the circulation and seroprevalence of ZIKV in LAC, owing to challenges in interpretation of serological tests that cross-react with other flaviviruses (e.g. dengue).^6,7^ In addition to the reduction in Zika cases there has also been a marked reduction in incidence of reported dengue and chikungunya cases in Brazil, meaning that the role of climatic, and other factors affecting mosquito density, or cross-immunity between arboviruses cannot be ruled out.

While the decline in ZIKV incidence is undoubtedly a positive development, it exposes clear gaps in our understanding of its natural history and epidemiology, which limit our ability to plan for, detect, and respond to future epidemics. The short duration of the epidemic and the long lead time needed to investigate comparatively rare congenital impacts has meant maternal cohort studies, in particular, may be statistically underpowered to assess relative risk and factors associated with ZIKV-related adverse infant outcomes.^8^ The evaluation of the safety and efficacy of ZIKV vaccine candidates^9^ are now also faced with an increasingly scarce number of sites with sufficient ZIKV incidence.^10,11^

There is an urgent need to predict which areas in LAC remain at risk of transmission in the near future, and estimate the trajectory of the epidemic. Projections can help public health policymakers plan surveillance and control activities, particularly in areas where disease persists. They can also be used by researchers, especially those in vaccine and drug development to update sample size calculations for ongoing studies to reflect predicted incidence within the time-window of planned trials. The findings identified from a continental analysis of ZIKV in LAC may be useful should ZIKV emerge in other settings, such as quantifying the spatial patterns of spread and impact of seasonality on incidence.

Several mathematical and computational modelling approaches have been developed to forecast continental-level ZIKV transmission ^5,11–14^. The focus has largely been on estimating which areas are likely to experience epidemic growth. It is apparent from the incidence in 2017 that many countries no longer reported an increasing incidence of cases. Due to either data unavailability or inaccuracies in the reported number of Zika cases in each country at the time of analysis, such approaches have either not used incidence data at all, ^15–17^ fit models to data on other arboviruses or used selected Zika-related incidence data from particular countries ^5,12,13,18–21^ to calibrate their models. Additionally, only a small number of studies have validated their model findings, either through comparison to serological surveys or comparing model outputs to incidence data not used within model fitting. ^13,19–21^ Considerably more data are now available across LAC and spanning multiple arboviral transmission seasons. This provides a valuable opportunity to examine the nature of ZIKV transmission and the importance of connectivity and seasonality in assessing ZIKV persistence in specific locations throughout LAC.

In this article, we apply a dynamic spatial model of ZIKV transmission in 91 major cities across LAC and fit the model to the latest data from 35 countries in LAC. We test several models to account for human mobility to better understand the impact of human movements on the emergence of ZIKV. The model was validated using a 10-fold cross-validation comparison to the data. We use the fitted model to quantify the expected number of cases likely to be observed in 2018 and identify cities likely to remain at greatest risk.

## Methods

### Zika case data from Latin America & the Caribbean

The weekly number of confirmed and suspected Zika cases within each country is reported to the Pan American Health Organization. This analysis makes use of the weekly incidence of Zika cases in 35 countries, from January 2015 to August 2017 (see Supporting Information S1). State-level ZIKV incidence data was available for Brazil and Mexico.^22^ Confirmed cases are typically identified through a positive real-time reverse polymerase chain reaction blood test using ZIKV-specific RNA primers. Suspected cases are based on the presence of pruritic (itchy) maculopapular rash together with two or more symptoms such as fever, or polyarthralgia (multiple joint pains), or periarticular oedema (joint swelling), or conjunctival hyperaemia (eye blood vessel dilation) without secretion and itch.^23,24^ Confirmed and suspected cases were included in this analysis because ZIKV detection may have low sensitivity due to a narrow window of viraemia and many samples, particularly from the earlier phase of the epidemic remain untested due to laboratories being overloaded during the epidemic.^24^ Inclusion of suspected cases in the analysis may reduce specificity due to the non-specific clinical manifestations of ZIKV and similar circulating arboviruses, including dengue. The reporting of ZIKV cases will vary considerably between settings and is thought to depend on the arbovirus surveillance system already in place, additional surveillance specifically established for ZIKV and other viruses and the likelihood of an individual self-reporting with symptoms consistent with ZIKV infection.

### A mathematical model of ZIKV infection

A deterministic meta-population model was used for ZIKV transmission between major cities in the LAC region. Cities with a population larger than 750,000 and large Caribbean islands were included in the model. In total we considered 91 cities. We extracted population sizes using the UN estimates from 2015.^25^ Migration between cities was modelled assuming several scenarios: i) a gravity model with no exponential terms; ii) a gravity model with estimated exponential terms; iii) a radiation model; iv) a data-driven approach based on flight data; and v) a model of local radiation and flight movements. Gravity models assume that movement between cities is highest when located near each other and when both cities are large. Radiation models assume that movement between cities are affected by the size of the population in a circle between the cities (see Supporting Information S2 for further information).

Within each city, individuals were classified by their infection status: susceptible, pre-infectious, infectious or recovered from ZIKV infection (Figure 1). Upon infection, individuals were assumed to be pre-infectious for an average of 5 days and then infectious for a subsequent 20 days.^26,27^ Immunity was assumed to be life-long and no cross-protection against other flaviviruses was assumed. The main vectors for ZIKV in LAC are thought to be *Aedes aegypti*, whilst *Aedes albopictus* and other species were thought to play a minor role in transmission.^28^ The seasonality and scale of ZIKV transmission was assumed to be specific to each city and dependent across cities, using a vector capacity modelling approach to estimate transmission intensity of the vectors and an environmental niche model to estimate vector relative abundance.^29–31^ In this approach, we model the probability that ZIKV may transmit for each day of the year, and feed this time-varying probability into the mathematical model. We estimate the time-varying reproduction number (R_0_(t)), defined the average number of secondary infections that result from one infected person within a totally susceptible population, that value varies in time due to the seasonality in vector capacity within each city. The seasonality curves are summarised using the average number of days per year where R_0_(t) was greater than 1, and the mean value of R_0_(t).

**Figure 1.**
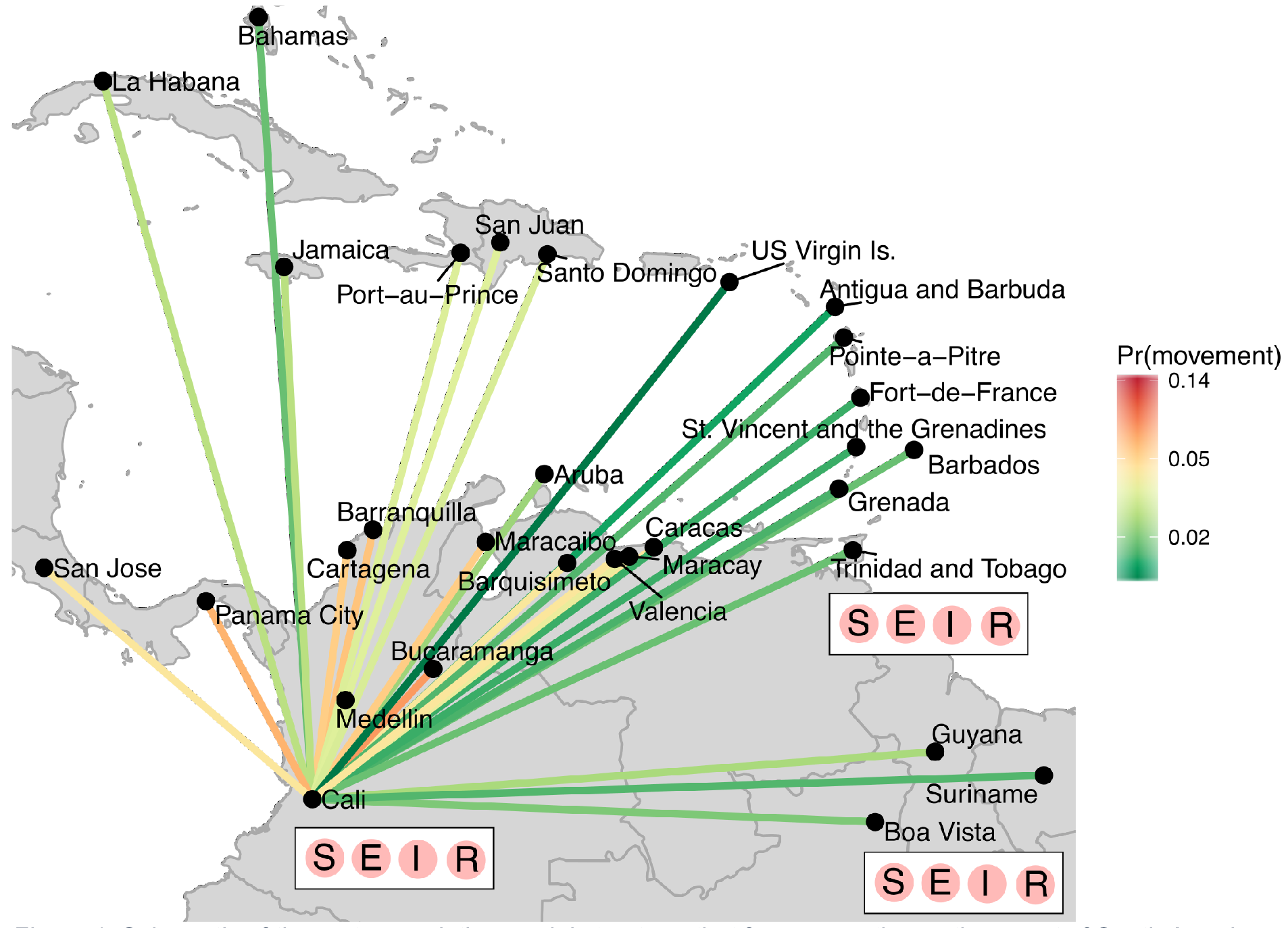
*Schematic of the meta-population model structure, that focuses on the northern part of South America and the Caribbean islands. Each city consists of individuals who are assumed to be susceptible (S), pre-infectious (E), infectious (I) or recovered (R) from ZIKV infection. Movement of pre-infectious individuals between cities is modelled assuming different population flows, where a gravity model is illustrated. Movements to cities outside of the plotted area are not illustrated.*

Owing to the difficulties in ZIKV disease surveillance,^23^ the weekly incidence of reported cases was unlikely to reflect the true incidence in each setting and we did not fit the model to weekly incidence data. We therefore used summary statistics in the model fitting procedure, focussing on the timing of the peak in incidence and whether the annual incidence was above 1 case per 100,000 in each country. The timing of the peak in outbreaks has been previously shown to be a useful summary statistic for epidemic dynamics,^32,33^ and preliminary analysis illustrated that annual incidence had a good discriminatory power for the estimating parameters of the model. Although surveillance quality varies between settings the timing of the reported peak within countries is less sensitive to systematic error. A sensitivity analysis confirmed that only a small number of observations were susceptible to large changes in surveillance prior to April 2016 and after January 2017, making the reported timing of the peak robust to changes in surveillance (Supporting Information S3).

The model estimate of new infections within each city was aggregated to the country or state level (for Brazil and Mexico) and scaled to ZIKV cases, enabling comparisons with the available data. The maximal value of R_0_(t) and the best-fitting migration model (including the maximal leaving rate from cities) were estimated in the model fitting procedure. Parameters were estimated using approximate Bayesian computation-sequential Monte Carlo (ABC-SMC) methods.^34^ ABC methods use summary statistics to estimate model parameters from qualitative epidemic characteristics. The sequential procedure of ABC-SMC means that each model of human mobility could be treated as a parameter. The prior and posterior distributions of selecting each model was used to estimate Bayes Factors to determine the evidence in favour of one model over another. Multiple parameter sets with equivalent fit were produced during the model fitting, and were used to provide the mean and 95% credible intervals (CI) of parameter estimates, numbers infected between 2015–2017, timing of the peak in the epidemic and projections of the numbers of ZIKV cases in 2018. The distribution of the timing of the peak was compared to the data using Bayesian posterior checks. The values correspond to probability that the data take a value less than or equal to the cumulative distribution function of the model, and values between 0.01–0.99 can be interpreted as evidence that the data and model estimate come from the same distribution. For each country the time series of reported cases were compared to the normalised model incidence. We compare the total number of reported cases to the estimated median (and 95% CI) number of infections to estimate the country-specific probability of reporting a case per infection.

To validate the parameter estimates and model output a cross-validation approach was used. The data was split into ten randomly allocated groups by country, each group was sequentially excluded from the parameter estimation procedure and the peak timing of the out-of-sample parameter estimates were compared to the data. The 95% CI of the cross-validated estimates were compared to the within-sample peak estimates. For the 2018 projections we reported the median number of cases, accounting for the estimated reporting rate and uncertainty in model output. The 95% prediction interval had a variance equal to the sum of the variance of the model prediction and the variance of the expected value assuming a Poisson distribution. Comparison of 2018 predictions to data were not possible as data from affected countries have not been made publicly available (as of 2 May 2018).

To date, there is no evidence to suggest sexual transmission contributes to population transmission,^35,36^ so this transmission route was not considered in the model. Due to current unexplained variability,^37^ we do not project the expected numbers of neonatal malformations or neurological disorders, such as microcephaly, associated with ZIKV infection.

## Results

A gravity model, which assumes migration scales with large populations that are closely located to one another, provided the best fit the data (Table 1). We identified substantial spatial heterogeneity in transmission (country summaries are provided in Table 2); the average estimated value of R_0_ was 1.81 (95% CI 1.74–1.87) and the average number of days per year where R_0_(t)>1 was 253 days (95% CI 250–256 days). The average number of days where R_0_(t)>1 varied from 116 days days (Costa Rica) to almost year-round transmission (several cities within Brazil (Belem & Salvador) and Colombia (Medellin & Cali), and Aruba and Curacao Islands). The maximal estimate of R_0_(t) was often higher within cities that also reported a longer window of transmission with R_0_(t)>1. However, several cities (including Boa Vista, Aracaju and Natal in Brazil) were estimated to have maximal R_0_(t) values above 2.5 with a relatively small window of transmission within the year.

**Table 1.** *Summary of the evidence for each population movement model tested on the Zika data. The prior and posterior probabilities were estimated using the ABC-SMC procedure (see SM for further details)*.

| Model of population movements | Gravity (simple) – $M_1$ | Gravity (exponential terms included) – $M_2$ | Radiation– $M_3$ | Flight data– $M_4$ | Combination of flight & radiation– $M_5$ |
| --- | --- | --- | --- | --- | --- |
| Prior probability ($\pi(m_y)$) | 0.232 | 0.246 | 0.224 | 0.052 | 0.092 |
| Posterior probability ($P(m_y\vert x)$) | 0.001 | 0.344 | 0.001 | 0.001 | 0.001 |
| Bayes Factor | 0.003 | 1 | 0.002 | 0.001 | 0.001 |
| Evidence for alternative model (and against model $m_y$) | Very weak evidence of fitting data | Model has best evidence of fitting data | Very weak evidence of fitting data | Very weak evidence of fitting data | Very weak evidence of fitting data |

Despite the emergence of the ZIKV epidemic in early 2015 in north-eastern Brazil, the incidence of cases remained relatively low in 2015 (Figure 2D and S5 for plots of Brazilian States). All countries that reported cases in 2015 (Brazil, Colombia, Guatemala, Honduras, Paraguay, Suriname, Cuba, El Salvador, Mexico, and Venezuela) continued to report cases in 2016 and 2017, except for Cuba. For most countries, the largest number of cases were reported in 2016. Belize, Colombia, French Guiana, Honduras, Suriname and several Caribbean islands reported more than 2 cases per 1,000 population in 2016. For 28 of the 35 countries in the analysis, the peak in reported disease incidence occurred in 2016. Five countries reported a peak in 2017 and Cuba reported a peak in July 2015 (Figure 2C).

**Figure 2.**
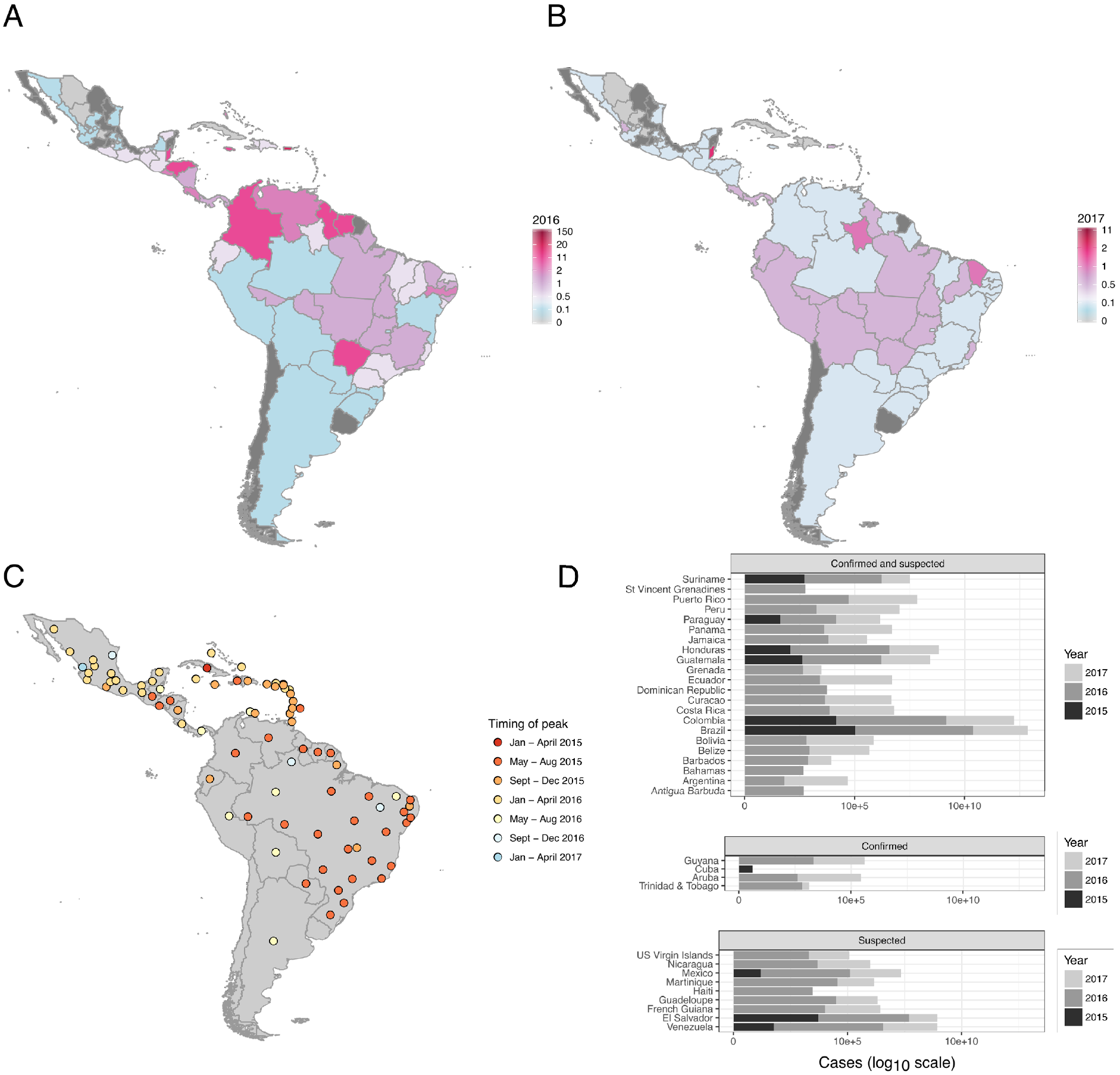
*Reported Zika incidence (cases per 1,000) within Latin America for A) 2016 and B) 2017, C) timing of peak incidence and D) total number of cases reported for each country for each calendar year (on a log 10 scale)), according to the case classifications submitted by each country.*

The estimated incidence of ZIKV infections (median and 95% CI) were compared to the reported data to estimate the country specific reporting rate. The average probability of an infection being reported as a case was 3.9% (95% CI 2.3–8.1%) and this rate was lower within countries that only reported confirmed cases (4 countries) than those who reported both confirmed and suspected cases (22 countries) (Table 2). Costa Rica, French Guiana and the US Virgin Islands were estimated to have a reporting rate above 20%. A comparison of the time-series of reported cases was compared to the model estimates of incidence (Figure 3). For all countries an epidemic was likely to have begun by Dec 2015 to Mar 2016 (otherwise known as the first phase). The relative scale of the epidemic in the first phase compared to late 2016 (the second phase) varied by country. For many countries the epidemic was estimated to be larger during the first phase (such as Argentina, Bolivia, Ecuador, Paraguay). For simulations in Antigua and Barbuda, Mexico and Venezuela the epidemic during the second phase had a higher incidence than the first phase. A small number of countries were estimated to have experienced only one epidemic season: Belize, Honduras, El Salvador and most Caribbean Islands. The difference in the timing of the peak between data and model was measured using Bayesian posterior checks where there was a non-significant difference between the model and data for 11 countries, and the distribution was over-dispersed (Figure 4A and 4B). There was a significant correlation (p=0.035) between the reported and estimated peak in the country epidemics (Figure 4C). The estimated peak in cross-validated simulations were correlated (p<0.001) with the model fit, although the 95% CI were wider (Figure 4D).

**Figure 3.**
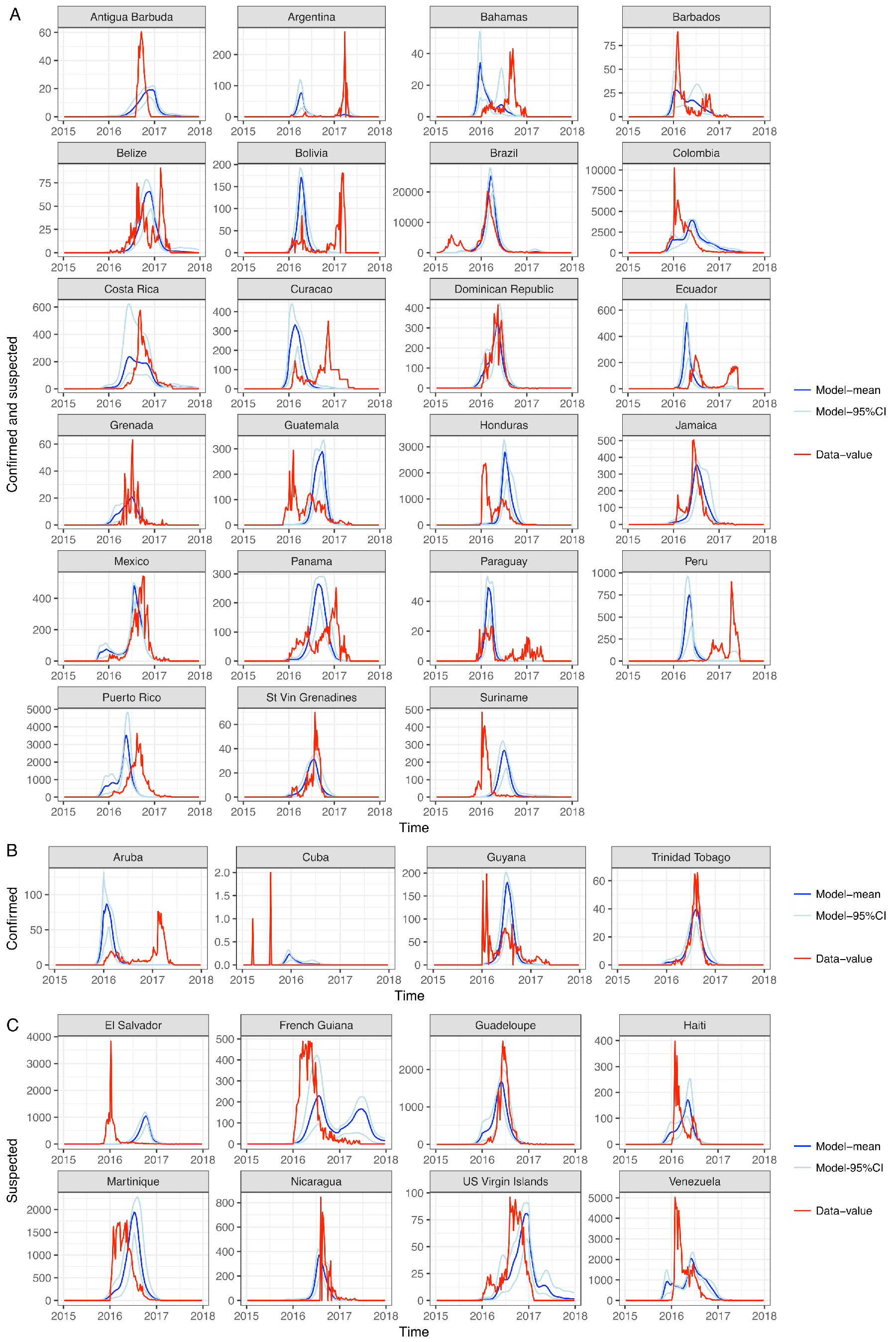
*Comparisons of the time-series data and normalised model output for all Latin American countries. The countries are ordered by the type of surveillance data available: Confirmed and suspected, Confirmed, and Suspected cases. To enable comparison we estimate the symptomatic reporting rate for each country and multiply this by the model estimate of the number of infections.*

**Figure 4.**
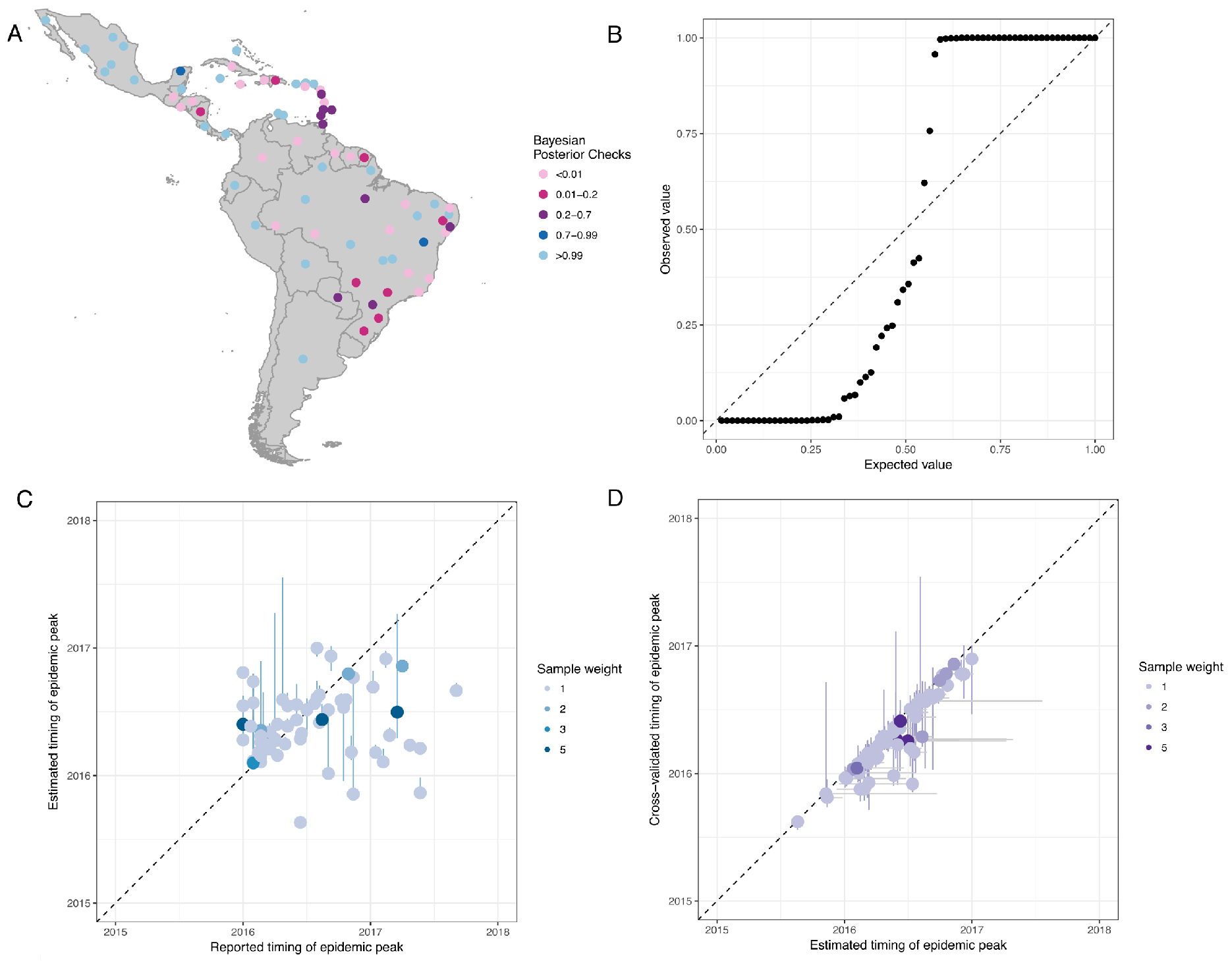
*Comparisons of observed and model fit for ZIKV peak incidence in the 31 countries in Latin America. A) Bayesian posterior checks that the estimated peak timing are consistent with the data; values between 0.01–0.99 indicate that the model and data are from the same distribution, B) Quantile plot of the Bayesian posterior probabilities C) Comparison of the observed timing of the peak and estimated timing of the peak (with 95% CI) D) Comparison of the estimated timing of the peak and the cross-validated estimates of peak timing (with 95% CI on the horizontal and vertical).*

**Table 2.**
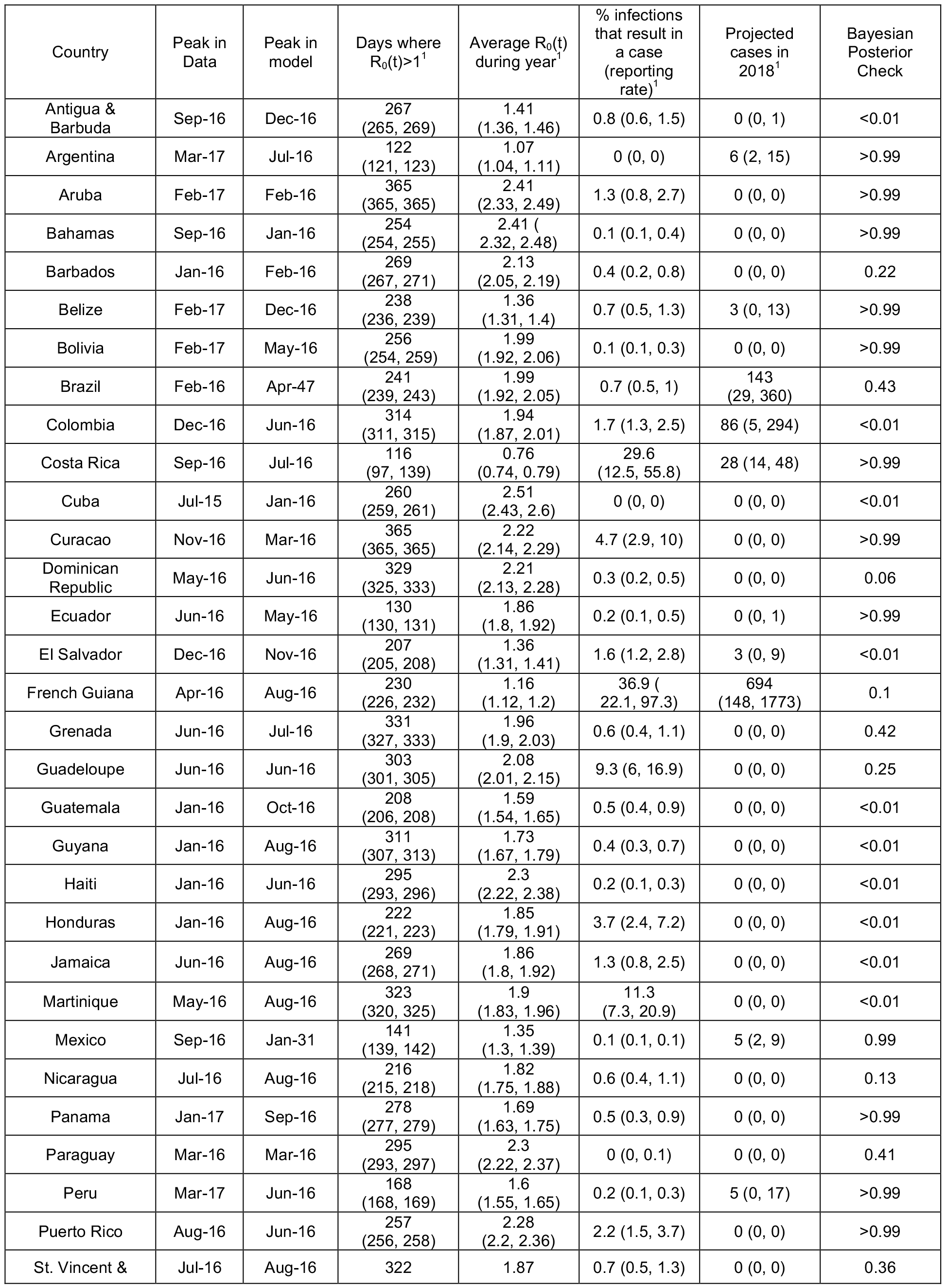
***Reported and estimated statistics for ZIKV in Latin America and the Caribbean.** Reported timing of the peak of ZIKV cases; the model estimate of the peak in ZIKV cases; the estimated number of days each year where R_0_>1; the average value of R_0_ throughout the year, the estimated reporting rate of ZIKV cases and the estimated number of ZIKV cases in 2018.*

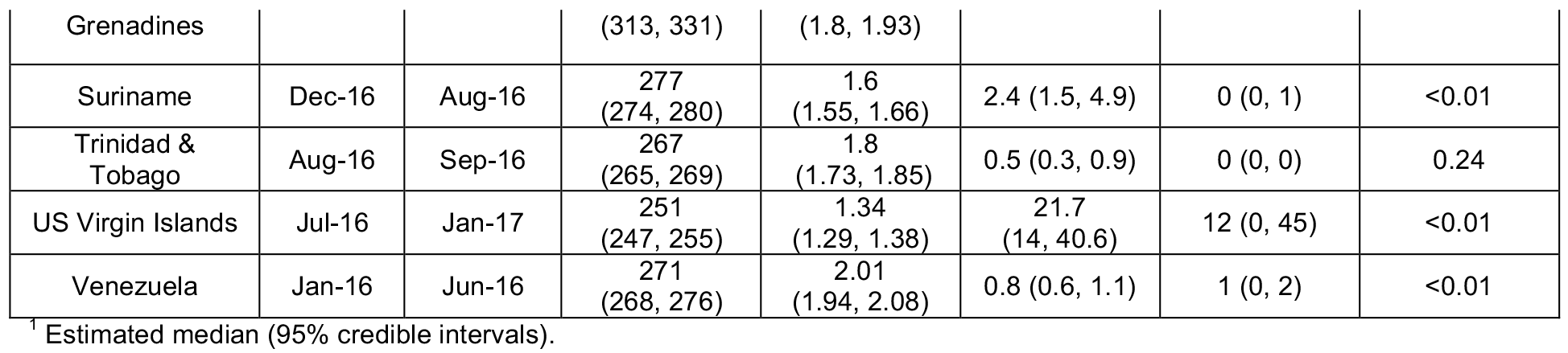

Projections for 2018 suggest a low incidence of Zika cases in most cities considered in the analysis (Figure 5, and Table 2). When accounting for the country-specific case reporting rate, the median number of cases was typically less than 20 in most settings. However, French Guiana was predicted to have between 148–1773 cases, owing to a larger pool of susceptible individuals than in other settings. Populated States within Brazil, such as Santa Carina and Säo Paulo were projected to have more than 5 cases, and cases were predicted to occur within Medellin (Colombia) and San Jose (Costa Rica). The majority of Caribbean countries were predicted to have no cases in 2018. For all cities the incidence of cases in 2018 will be lower than 2017. In Colombia, the projected time series of cases for specific cities illustrate a negligible incidence in 2018, but Medellin was expected to experience the end of the epidemic in 2018 (Figure 5C).

**Figure 5.**
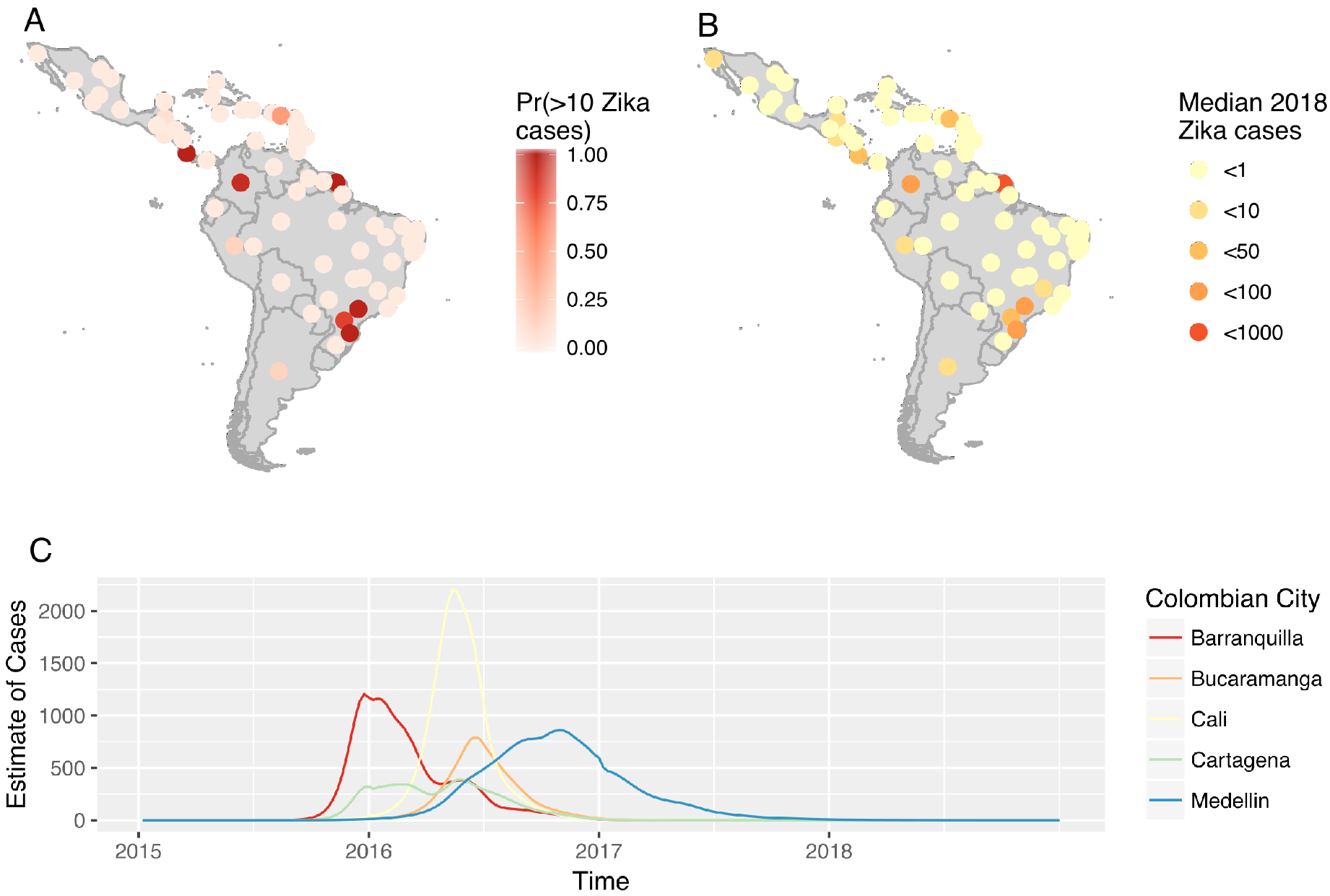
*A) The estimated probability of Zika cases in each country (and States in Brazil and Mexico) Probability of more than 10 cases B) Median estimate of Zika cases in 2018 C) The estimated time series of Zika cases within 5 major cities of Colombia.*

## Discussion

The spread of ZIKV across the LAC region in 2015–2017 has resulted in considerable disease burden, particularly in the children of mothers infected during pregnancy. Both the reported incidence of cases and modelling results from this study suggest that the transmission of ZIKV had continued until herd immunity was reached, despite major efforts to limit its spread through vector control. Whilst the reported and projected reduction in ZIKV cases is undoubtedly good news for affected communities, it is only because substantial numbers of individuals have already been infected. Therefore, it remains vital to maintain surveillance for congenital and developmental abnormalities and provide long-term care for affected people and families.^38^

The aim of this analysis was to assess if cities in LAC were likely to experience ZIKV cases in 2018, to support resource planning and trials. Our modelling results suggest a very low incidence in 2018. This analysis supports the findings of previous mathematical models of ZIKV. ^5,11,13,14^ In addition, our study provides estimates of incidence and risk for specific cities, estimates of case reporting rates, incorporates parameter uncertainty, includes out-of-sample validation of the model estimates and uses more data than other modelling studies as we incorporate ZIKV case reports alongside ecological data to determine city-specific epidemic trajectories and seasonality curves.

We fitted the model to the timing of the peak in ZIKV cases and then compare the time series of expected cases to reported cases and found a good fit in many countries. We assumed that large cities both drive the spread of Zika and are responsible for the majority of cases. As *Ae. aegypti* is a largely urban dwelling mosquito and that arboviral diseases have been observed to be spread by movement of infected humans,^39,40^ this assumption is likely to be valid. However, while we predict the outbreak to largely be over in these large cities, smaller more remote cities and peri-urban areas may still have susceptible individuals and experience cases. Should additional sub-national data on the timing of the peak become available, the model fitting and projections can easily be updated. Case reporting rates indicate a lower rate within countries that report only confirmed cases, and the rates within Brazil, El Salvador, Martinique, Puerto Rico, and Suriname align well with other estimates measured using alternative methods.^21,41,42^

Despite the short-comings in the available data, we present the most up-to-date and robust predictions of Zika incidence in 2018. As the projected incidence is consistently low across all model runs, this finding is quite robust to the variability accounted for in the model. Validation of these findings are necessary through multi-site population representative seroprevalence surveys across LAC to monitor seroconversion to ZIKV, such as in Netto et al..^19^ Reporting of cases within LAC has reduced markedly since the downgrading of ZIKV from a public health emergency of international concern to an ongoing public health challenge (in November 2018).^43^ Consequently, it remains difficult to compare these projections to incidence data for 2018.

This research has highlighted that within LAC the spread of ZIKV was better represented by a gravity model than flight movements. This may seem surprising as flight data are cited as a source of emerging infections, such as ZIKV.^44^ But cars and public transportation are used for most journeys; and the movement of people impacts the spatial spread of vector-borne diseases.^38,45^ Perhaps for highly transmissible infectious diseases movements facilitated by flights are sufficient for predicting introduction of a pathogen in a new population, but this analysis suggests that triggering of a ZIKV outbreak may require more frequent exposure than air travel. The migration patterns assumed within each model are quite different in LAC (see Supporting Information), suggesting that models which have not tested the relative fit of each and use one alone could be prone to errors in estimated spread of ZIKV. In comparison to mobility modelling in North America, Europe and Africa, the mobility patterns in Latin America are not well quantified and require further study.

Major questions on the epidemiology of ZIKV remain unanswered.^7^ Whilst the impact of sexual transmission on ZIKV emergence is likely to be minimal,^35,46^ it may increase the magnitude of an epidemic^36^ and this would be difficult to test using the available surveillance data. There are large differences in the incidence of congenital Zika syndrome across LAC,^38^ with an epicentre reported within northeast Brazil, that remain largely unexplained. In particular, the analysis here suggests increased incidence of Zika throughout Brazil in 2016, but the expected increase in congenital malformations within newborns were not observed.^47^ This and other modelling studies suggest that ZIKV has been widespread, and the finding of geographically variable rates of congenital defects is discordant with the more consistent rates of ZIKV infection predicted by our model.

We have assumed that the time varying transmission rate of ZIKV is a function of environmental and vector suitability that has not been reduced by effective vector control. The impact of vector control has been largely unassessed, or where it has been assessed it has been found to be ineffective.^48,49^ Consequently, our findings are likely to be unaffected by the impact of vector control. Should effective wide-scale interventions be developed, the model can be used to assess the impact of proposed interventions. The mathematical model was deterministic in nature, and especially for projections may under-estimate the variability in the number of cases. Additionally, we do not include the impact of inter-annual variation in *Aedes aegypti* vector capacity, such as the 2015–2016 El Nino climate phenomenon, which has previously been shown to be positively associated with an increased incidence in 2016.^18^ Instead, we show that the peak incidence in 2016 was likely due to a low incidence of infection in 2015, that then resulted in optimal transmission in 2016, resulting in depletion of the susceptible population, thus limiting incidence in 2017 and 2018. If inter-annual variation in ZIKV transmission were incorporated into our model, it is likely that our incidence estimates for 2016 would increase, and the predicted incidence in subsequent years would further decrease.

## Conclusions

ZIKV has spread widely across LAC, affecting all cities during 2015–2017 leading to high population immunity against further infection, thereby limiting capacity for sustained ZIKV transmission. The seasonality in ZIKV transmission affected the rate of infection, but due to high connectivity between cities this had little impact on the eventual depletion of susceptible populations. Looking forward, incidence is expected to be low in 2018. This provides optimistic information for affected communities, but limits our ability to use prospective studies to better characterise the epidemiology of ZIKV. The continental-wide analysis illustrates much commonality between settings, such as the relative annual incidence, and the connectivity across LAC, but questions remain regarding the interpretation of the varied data for ZIKV. Ultimately, representative seroprevalence surveys will be most useful to understanding past spread and future risk of ZIKV epidemics in LAC.

## Abbreviations

CI: credible intervals
LAC: Latin America and the Caribbean
ZIKV: Zika virus

## Declarations

### Ethics approval

Institutional ethics approval was not sought because this is a retrospective study and the databases are anonymised and free of personally identifiable information.

### Availability of data and material

The models developed here and data are available at https://github.com/kath-o-reilly/Zika LAC-Outbreaks.

## Competing interests

The authors declare that they have no competing interests.

## Funding

This work was partially supported by the European Union’s Horizon 2020 Research and Innovation Programme under ZIKAlliance Grant Agreement no. 734548 and ZikaPLAN Grant Agreement No 734584. RL was funded by a Royal Society Dorothy Hodgkin Fellowship. AK was funded by a Sir Henry Dale Fellowship jointly funded by the Wellcome Trust and the Royal Society (grant Number 206250/Z/17/Z). OJB was funded by a Sir Henry Wellcome Fellowship funded by the Wellcome Trust (grant number 206471/Z/17/Z). SC was funded by the Laboratoire d’Excellence Integrative Biology of Emerging Infectious Diseases program (grant ANR-10-LABX-62-IBEID), the Models of Infectious Disease Agent Study of the National Institute of General Medical Sciences, the AXA Research Fund.

## Authors contributions

KO, RL, JE, PM, AK, RE, SC, OB and LY designed the study and KO, RL, DB, KK, OB and LY carried out the analysis. KO, RL, JE, PM, AK, RE, SF, DB, KK, MK, AW, LR, PB, EM, TJ, SC, OB and LY contributed to the data analysis and interpretation, writing of the report and approved it before submission.

## Acknowledgements

We thank partners within the ZikaPlan and Zika Alliance consortia who have provided comments and insight on previous versions of this work. We also greatly acknowledge the availability of publicly accessible data on ZIKV cases and other inputs of the model.

